# Dynamics of Florida milk production and total phosphate in Lake Okeechobee

**DOI:** 10.1101/2021.03.09.434552

**Authors:** Joseph Park, Erik Saberski, Erik Stabenau, George Sugihara

## Abstract

A central tenant of the Comprehensive Everglades Restoration Plan (CERP) is nutrient reduction to levels supportive of ecosystem health. A particular focus is phosphorus. We examine links between agricultural production and phosphorus loadings in the Everglades headwaters: Kissimmee River basin and Lake Okeechobee, considered an important source of water for restoration efforts. Over a span of 47 years we find strong correspondence between milk production in Florida and total phosphate in the lake, and, over the last decade, evidence that phosphorus in the lake may have initiated a long-anticipated decline in water column loading.

## Introduction

### Historical Perspective

Prior to the 19^th^ Century, the Florida Everglades consisted of 3 million acres of marsh draining the Kissimmee River Basin and Lake Okeechobee southward into Florida Bay. Water flowing into Lake Okeechobee came primarily from the Kissimmee River, meandering approximately 103 miles from Lake Kissimmee to Lake Okeechobee through a 1 to 2 mile-wide floodplain. The extensive floodplain kept nutrients at low concentrations throughout the system. As a result, addition of even small amounts of nutrients can significantly effect the structure and productivity of the native ecosystem [1].

Consistent with ideals of *manifest destiny*, efforts to “drain” the Everglades to produce arable lands were initiated in the late 19^th^ Century, and, in the 1950’s, the Kissimmee Flood Control project replaced the original meandering geometry with a channel consisting of straight-line segments [2, 3]. Completion of the project coincided with increased phosphorus loads to Lake Okeechobee from the transport of phosphorus-laden sediments [4, 5]. A comparative rendition of the pre-development and current systems is shown in figure 1.

**Fig 1.**
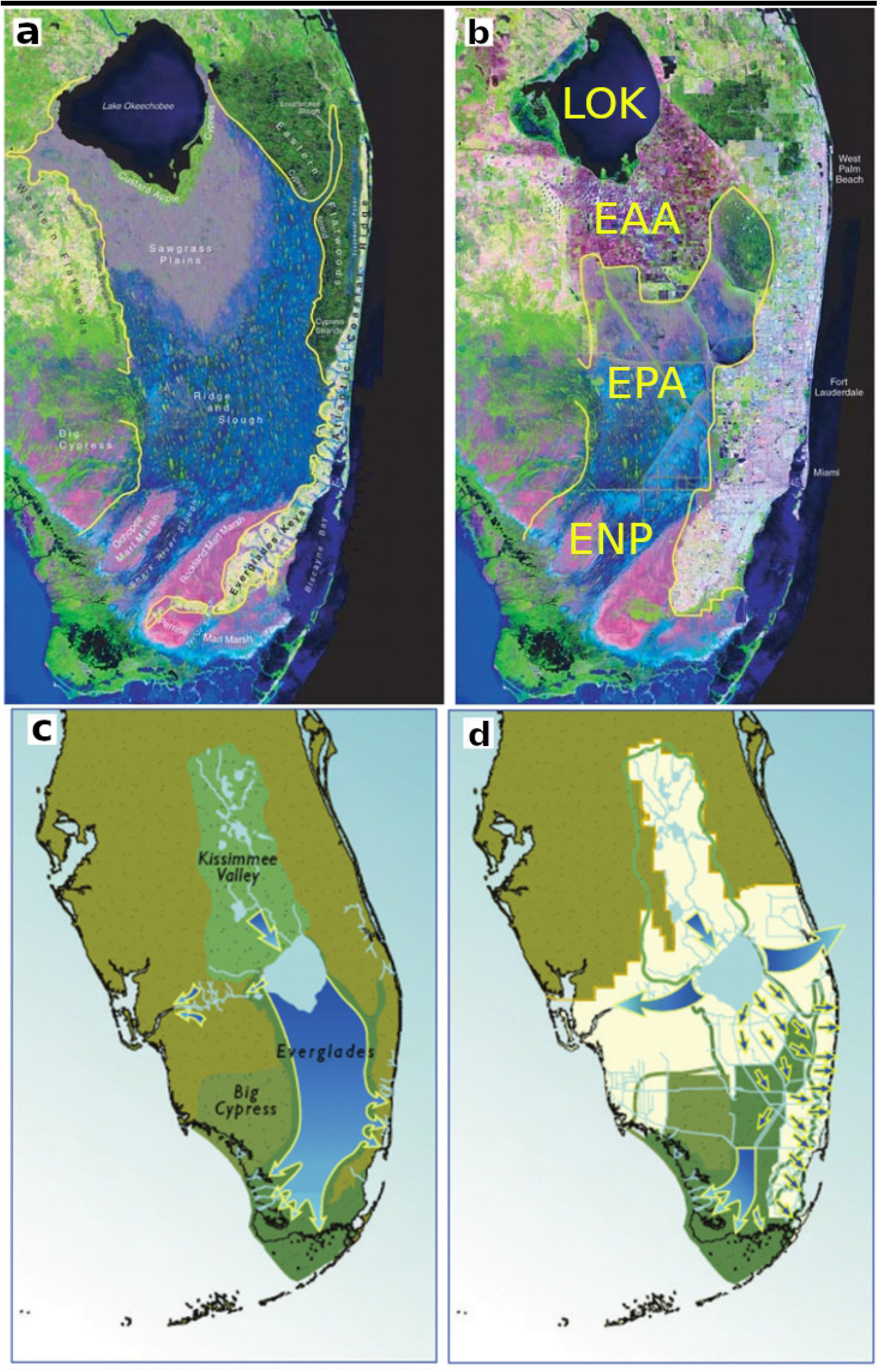
South Florida satellite image. a) Simulated South Florida satellite image circa 1850. b) Satellite image 1994. c) Pre-development hydrologic flow paths. d) Current flow paths. LOK -Lake Okeechobee. EAA -Everglades Agricultural Area. EPA -Everglades Protection Area. ENP -Everglades National Park.

The extensive spread of agriculture in the upstream drainage basins also contributed to this increased load. Phosphorus is added to uplands in fertilizers, organic solids (e.g., animal wastes, composts, crop residues), wastewater, and animal feeds. Some phosphorus is exported from the drainage basin as agricultural products, however, a significant amount accumulates in upland soils and sediments, and, a portion is then transported by surface flow to the lake [6].

Historically, cattle ranching was the main agricultural use of the watershed north of the lake, however, in the 1950s dairy farming increased eight-fold, with a corresponding increase in phosphorus exports from 250 to 2,000 metric tons per year [7]. In 2000, Florida enacted the Lake Okeechobee Protection Act (Chapter 00-103, Laws of Florida), mandating a comprehensive plan to reduce watershed phosphorus loading to meet a total maximum daily load (TMDL) of 105 metric tons (mt) per year of surface-water input by 2015.

### Contemporary Conditions & Restoration

Over the last two decades, many sources have been remediated, with appreciable declines in point source loads [7–15]. On-site monitoring at the farm level shows improvement, particularly for intensive land uses such as dairies where chemical treatment systems, stormwater management, and reuse systems have been implemented [16]. However, a large amount of legacy phosphorus remains throughout the watershed [17], and, owing to the significant variability in phosphorus dynamics, it is not clear that total phosphorus loading to the lake has significantly declined [18].

Because a substantial quantity of “new water” for the CERP will be delivered from Lake Okeechobee, water quality trends in the lake have important implications for Everglades restoration plans [18]. Although, at present, the nutrient-rich waters of Lake Okeechobee have limited effect on the downstream Everglades Protection Area (EPA), as roughly 4 percent of the lake’s outflow (on average) reaches the EPAs. Nutrients discharged from the Everglades Agricultural Area (EAA) and C-139 basins have been identified as the major sources impacting the downstream Everglades Protection Areas [3]. Source control strategies such as best management practices (BMPs) and stormwater treatment areas (STAs) have been implemented to reduce phosphorus loads from EAA basins to the Everglades Protection Area [19].

Specific to Lake Okeechobee, high phosphorus concentrations impact biota by altering the structure and functioning of the lake and downstream ecosystems. The overall increase of phosphorus loading has resulted in conversion of a phosphorus-limited system to a nitrogen-limited system, producing changes in the lake such as increased frequency of algal blooms and increased abundance of nitrogen-fixing cyanobacteria [20]. For example, during summer 2016, a large bloom of the cyanobacterium *Microcystis aeruginosa* occurred in Lake Okeechobee and subsequently in the St. Lucie Estuary, attributed to high nutrient levels supporting the growth of phytoplankton [18].

### Dynamical perspective

Conventional views find that total phosphorus loading to the lake has not significantly declined over the 1974-2017 period of record, despite the array of projects that have reduced phosphorous sources [12, 18]. However, there is significant variability across multiple timescales in the phosphorus data, with the potential to confound linear, block approaches of statistical interpretation. Here, we use two data-driven, nonlinear dynamical tools to examine time and cross variable dependence: Empirical Mode Decomposition (EMD) [21], and, Empirical Dynamic Modeling (EDM) [22].

EMD decomposes signals into scale-dependent modes termed intrinsic mode functions (IMF) without constraints of linearity or stationarity as presumed by Fourier, wavelet or Eigen decomposition. IMFs capture oscillatory modes, and, EMD residuals the nonlinear trends. Application of the Hilbert transform to IMFs provides time-dependent instantaneous frequency estimates, with the combination of EMD and IMF Hilbert spectra constituting the Hilbert-Huang transform (HHT). IMFs are particularly astute at isolating physically-meaningful dynamics. An introduction and review is found in reference [21].

EDM is a toolset for system analysis and forecasting based on multidimensional state-space representations of dynamical systems. EDM is not based on parametric presumptions, fitting statistics, or, specifying equations; providing a data-driven approach amenable to nonlinear dynamics [23]. A lucid introduction to EDM is provided by Chang et. al. [22].

We examine the data with EMD and EDM to reveal underlying dynamics and relationships between milk production and lake phosphorus. The synthesis of EMD and EDM has been termed empirical mode modeling (EMM), where EMD IMF’s are used to create physically relevant multivariable state spaces for EDM [24].

## Materials and methods

### Data

Milk production data are obtained from the United States Department of Agriculture (USDA) National Agricultural Statistics Service (NASS) database, reporting monthly milk production in Florida from January 1970 through June 2020 [25]. Data are shown in figure 2a.

**Fig 2.**
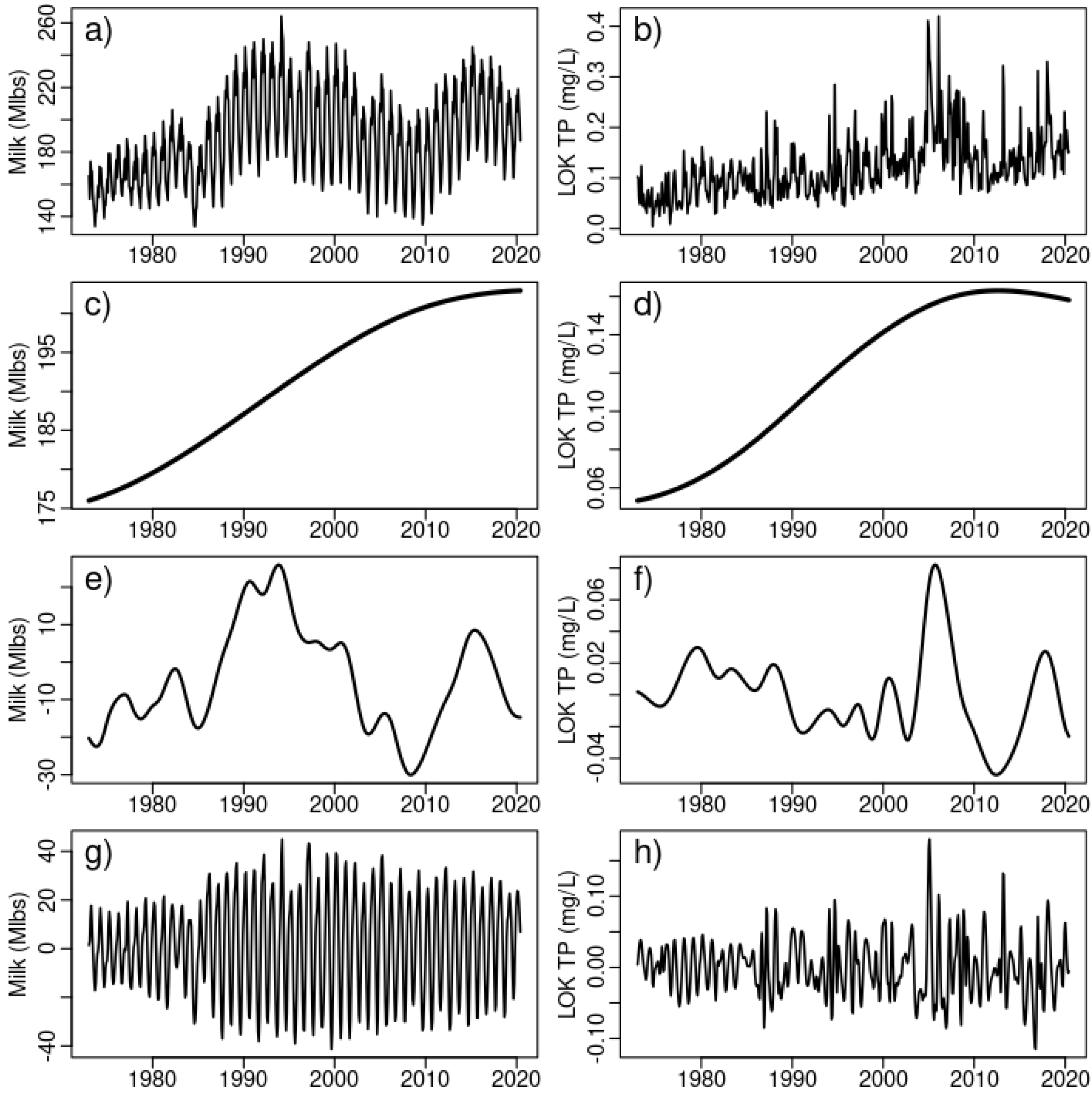
Data and empirical mode decompositions. a) Milk production b) LOK total phosphate c) Milk decadal d) Total phosphate decadal e) Milk interannual f) Total phosphate interannual g) Milk intra-annual h) Total phosphate intra-annual.

Phosphorus data for Lake Okeechobee is a 5 station average obtained from the South Florida Water Management District (SFWMD) DBHydro environmental database [26]. Raw data span the period December 11, 1972 to August 8, 2020. The time series is interpolated with a spline to monthly dates of the milk production data. The result is a data block of monthly milk production and interpolated total phosphate from January 1973 through June 2020 (figure 2a,b).

### 0.1 Empirical mode modeling

We use EMD [27] to decompose the time series into IMFs and associated nonlinear residuals (see figure 5 in Appendix Empirical mode decomposition). Interannual time series are created by summation of IMFs with interannual frequencies, for milk production IMFs 4,5,6, and IMFs 5,6,7 for phosphorus. Intra-annual time series consist of IMFs 2 and 3 for milk production, and, IMFs 2,3,4 of phosphorus.

We then use raw data and IMFs of interannual and intra-annual modes in an EDM convergent cross mapping (CCM) analysis [28]. CCM identifies potential causal links between state variables based on information shared between multidimensional embeddings of the variables [29]. CCM can be viewed as a dynamically-informed, fully nonlinear, analog to cross correlation. However, instead of reliance on temporal or cross variable *coincidence*, CCM is based on affine mappings of dynamical system states where CCM values indicate the cross variable predictability. Convergence of predictability as the information content and density of the state-space increase indicate shared dynamics and a measure of causality [29].

Since the data are autocorrelated (lag-1 correlations of 0.86 and 0.70 for milk and phosphorus respectively), and, exhibit seasonal dynamics, we use an exclusion radius of 12 points (months). That is, in the EDM state-space nearest neighbor search for the prediction at each time step, neighbors that are temporally within the exclusion radius of 12 points (months) are excluded from the prediction. This prevents any influence of autocorrelation or seasonality on the CCM information assessment.

To assess significance of the CCM results, we employ surrogate data samples created from the random phase method of Ebisuzaki [30]. We use N = 1000 surrogate time series of the milk or phosphorus data, and compute an EDM cross map strength for each surrogate and the original time series. For example, the original phosphorus time series is EDM cross mapped against 1000 surrogate milk time series created from Ebisuzaki spectral phase randomisation.

Given the cross map strength of variables *X* and *Y, ρ*_*XY*_, and, a vector of cross map strengths *ρ*_*XY*_*N* between *X* and N surrogates *Y*_*N*_, a p-value representing the probability of rejecting the null hypothesis that the cross map strength *ρ*_*XY*_ is not due to randomness, can be specified as:

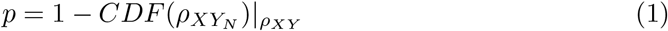

where 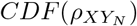 is the distribution function of the surrogate cross map strengths.

## Results

One of the most striking results can be seen in a relative comparison of milk production and lake phosphorus at different time scales. Figure 3 presents scaled (mean offset, standard deviation normalized) comparisons of the raw data, nonlinear trends, interannual, and, intra-annual modes. In the nonlinear trends of panel b) there is a remarkable coherence on the decadal time scale with relative changes in milk production and lake phosphorus nearly identical from the mid 1970’s through the early 2000’s. In the final decade, 2010-2020, there is flattening of overall milk production, and, a noticeable inflection towards phosphorus reduction. This may represent the long-anticipated decline in lake phosphorus loadings from surface water sources in response to best management practices and remediation efforts.

**Fig 3.**
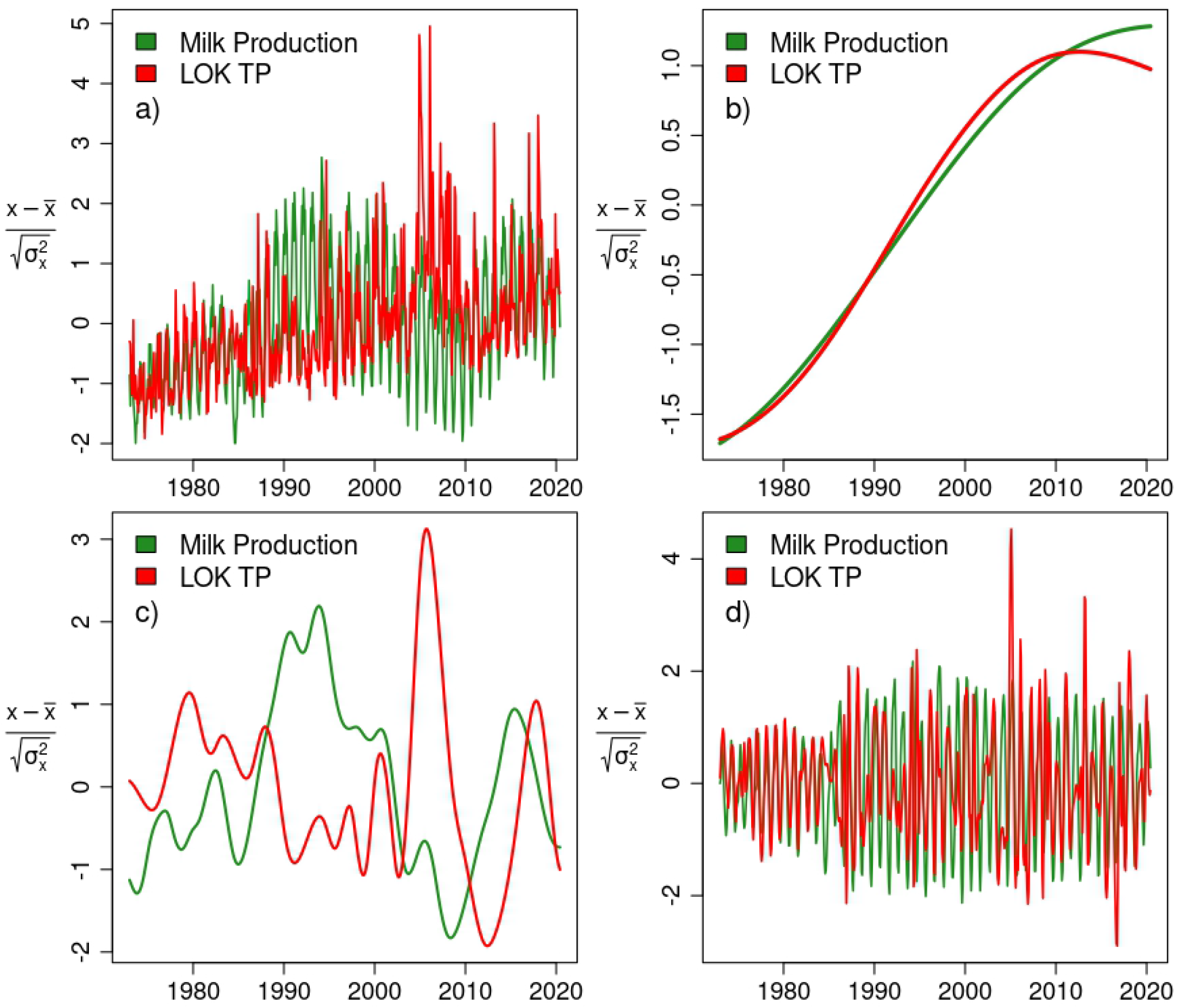
Empirical mode decompositions scaled. a) All timescales, b) decadal, c) interannual, d) intra-annual.

The interannual components provide no clear covariance, while the intra-annual comparison suggests strongly phase-locked dynamics over most years. To explore cross variable dependence, we use CCM.

### Convergent Cross Mapping

Convergent cross mapping assesses the extent to which states of variable *X* can be predicted from variable *Y*. If predictability using the entire time series (the full *library* of states) is significant, and, if predictability increases and converges as the state-space provides improved representations of the dynamics with increasing library size, it indicates shared dynamics and a causal link [29]. Figure 4 shows CCM results applied to milk production and phosphorus at different time scales. We note that the nomenclature for cross mapping is X:Y, indicating that X is used to predict states of Y. CCM tests for causation by measuring the extent to which Y can reliably estimate states of X. This happens only if X is causally influencing Y. This means that links for Y causing X are denoted X:Y.

**Fig 4.**
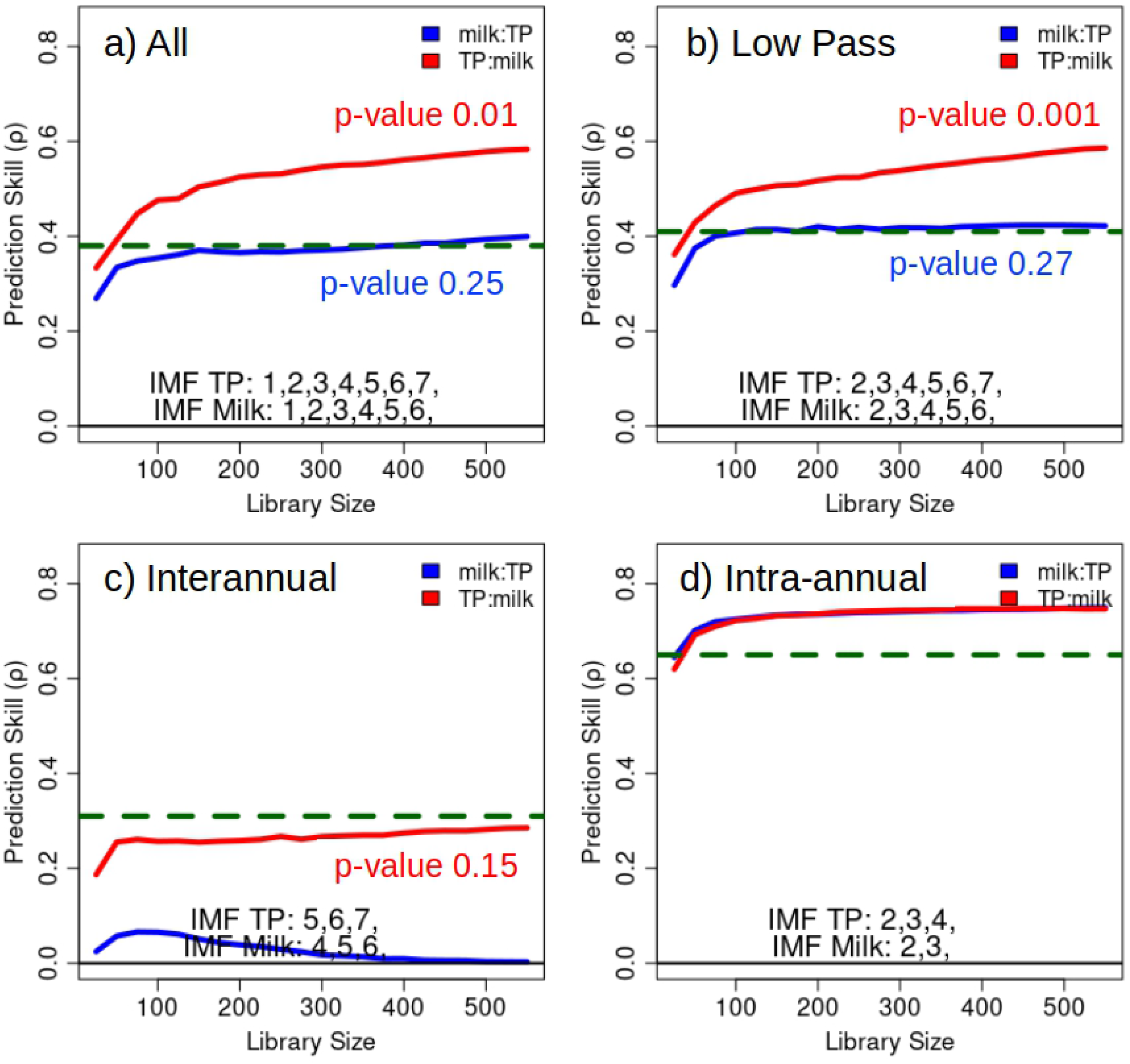
Convergent cross mapping (CCM) of milk and lake phosphorus. Dashed horizontal lines show linear cross correlation. a) All timescales, b) Low pass filter, c) interannual, d) intra-annual.

**Fig 5.**
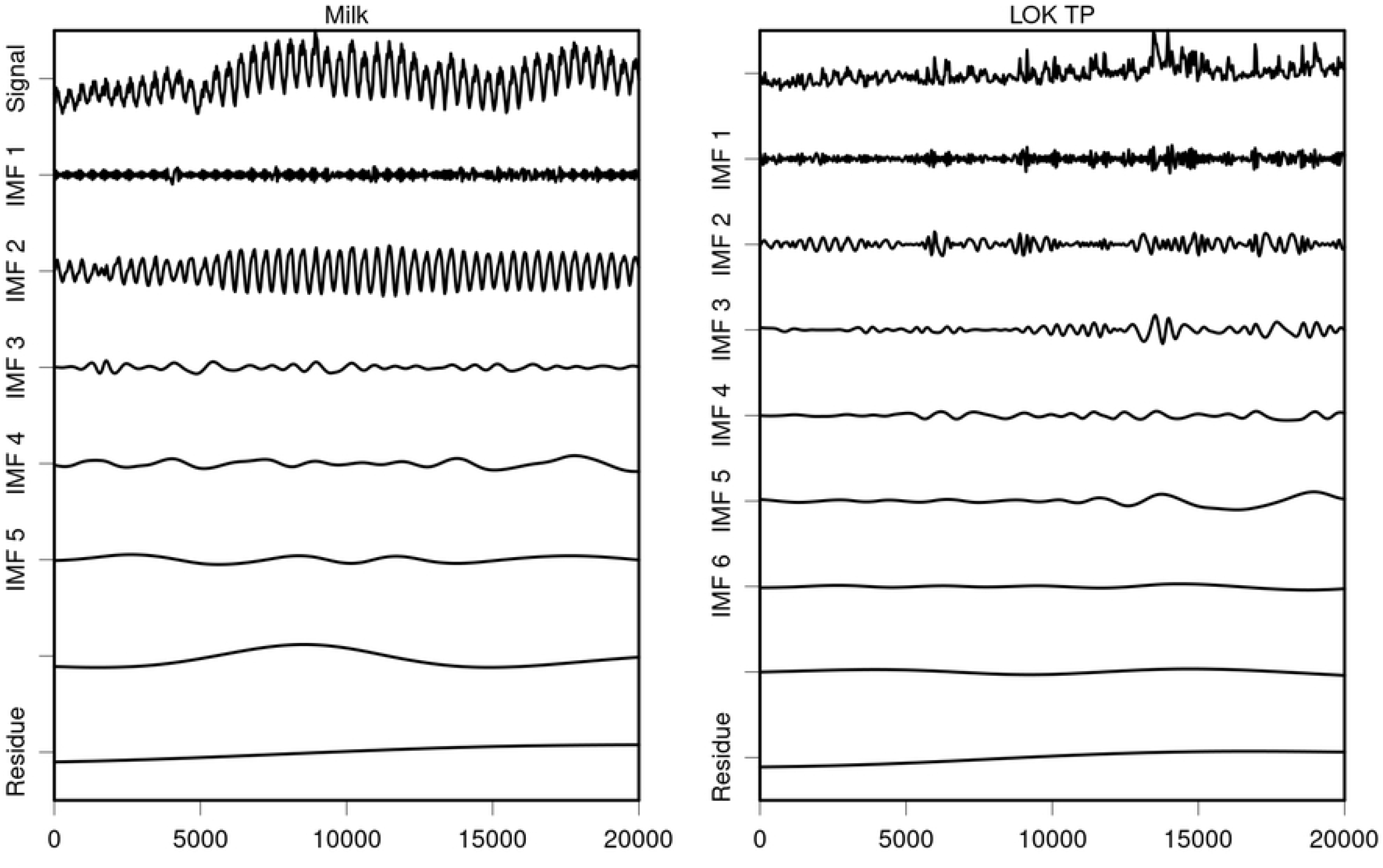
Empirical mode decompositions of milk production and lake phosphorus time series.

The top row of figure 4 plots CCM with all time scales a), and, with a low pass filter of the respective time series by removal of the highest frequency IMF b). Here, we find evidence that milk production can be considered a causal driver of lake phosphorus when all time scales are included in the system dynamics, and, slightly clearer evidence when high frequency noise is removed.

Panel c) indicates no viable link between milk production and lake phosphorus on interannual (ENSO) time scales. Appendix El-Niño Southern Oscillation and milk production provides additional details on the lack of a interannual link associated with ENSO.

The intra-annual, seasonal metric indicates strong coupling, but fails to resolve a directional attribute to the state predictions. This suggests a coupled, or common external driver on seasonal time scales.

## Conclusion

Decades of prosperity and attendant agricultural productivity have indelibly altered the land-use and ecosystems of central and south Florida. The addition of nutrient concentrations at order-of-magnitudes above pre-development levels have created a large reservoir of phosphorus in Lake Okeechobee. The waters of the lake are an important source of water supply for the Comprehensive Everglades Restoration Plan, but strict phosphorus limits must be met for this water to be admitted into the Everglades. Impressive phosphorus remediation efforts have been undertaken this Century, including restoration of natural flow paths to portions of the Kissimmee River [31], however, owing to the large amount of legacy phosphorus and complex dynamics of phosphorus monitoring, traditional data processing has not identified a decline of loading to the lake. Using tools from nonlinear dynamical systems analysis, we find evidence of a reduction in mean phosphorus loading to the lake over the last decade. Continued monitoring will reveal whether this reflects the long-anticipated secular trend in phosphorus reduction, or, a temporary decline.

Additionally, we find that when data are viewed across all time scales, there is an apparent causal link between milk production and phosphorus loading in the lake. This verifies the importance of continued remediation and source control efforts to mitigate phosphorus runoff.

## Supporting information

### Empirical mode decomposition

Fig. 5 shows the empirical mode decompositions of the milk production and lake phosphorus time series.

### El-Niño Southern Oscillation and milk production

To assess links between interannual dynamics of milk production and exogenous forcing, we explore the hypothesis that positive ENSO phases, El-Niño conditions, can result in increased milk productivity. The hypothesis is based on two underlying facts: First, that El-Niño conditions tend to produce cooler and wetter conditions in Florida [32], and second, a negative linear relationship between Holstien milk production and mean daily temperature [33]. That is, Holstien milk production increases as temperatures moderate from 82°F to 72°F. We therefore expect a positive relationship between El-Niño conditions and milk production. To quantify the state of ENSO, we use the Multivariate ENSO Index (MEI) [34].

Interannual components of milk production and MEI are shown in figure 6, where visually, there seems to be some correlation between milk production and MEI.

**Fig 6.**
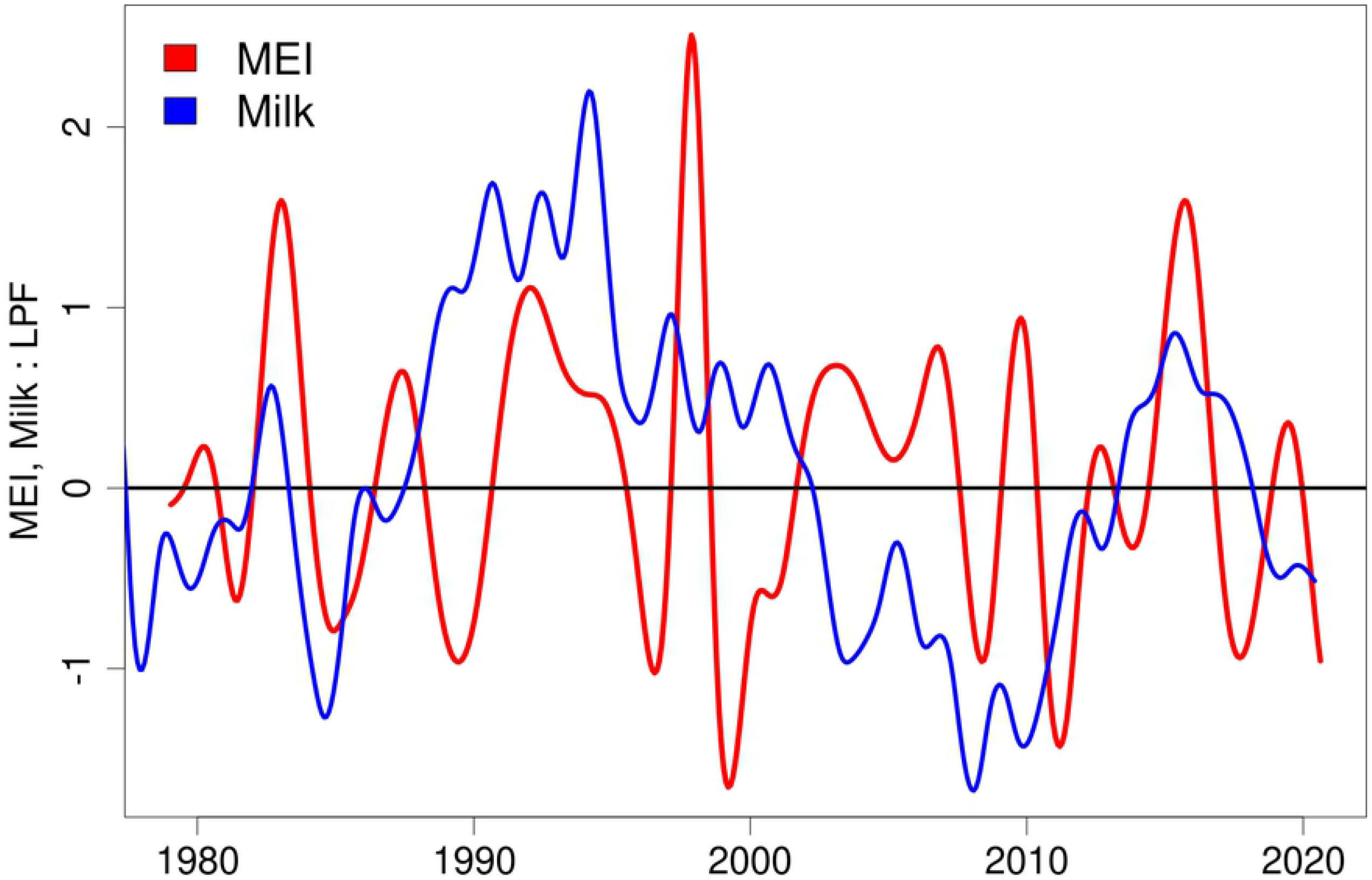
Interannual components of milk production and the Multivariate ENSO Index (MEI).

We assess links between these time series using CCM with results shown in figure 7, indicating no significant link between ENSO state and milk production. Interestingly, the linear correlation is weak, but statistically significant with a p-value of 0.0019. In contrast, the CCM analysis finds a p-value of 0.32, indicating an insignificant relationship. The inappropriateness of a linear model is clear when time series are viewed in a scatter plot, figure 8, clearly indicating nonlinear state dependence and a lack of dependence between MEI and milk production if state dependence is ignored.

**Fig 7.**
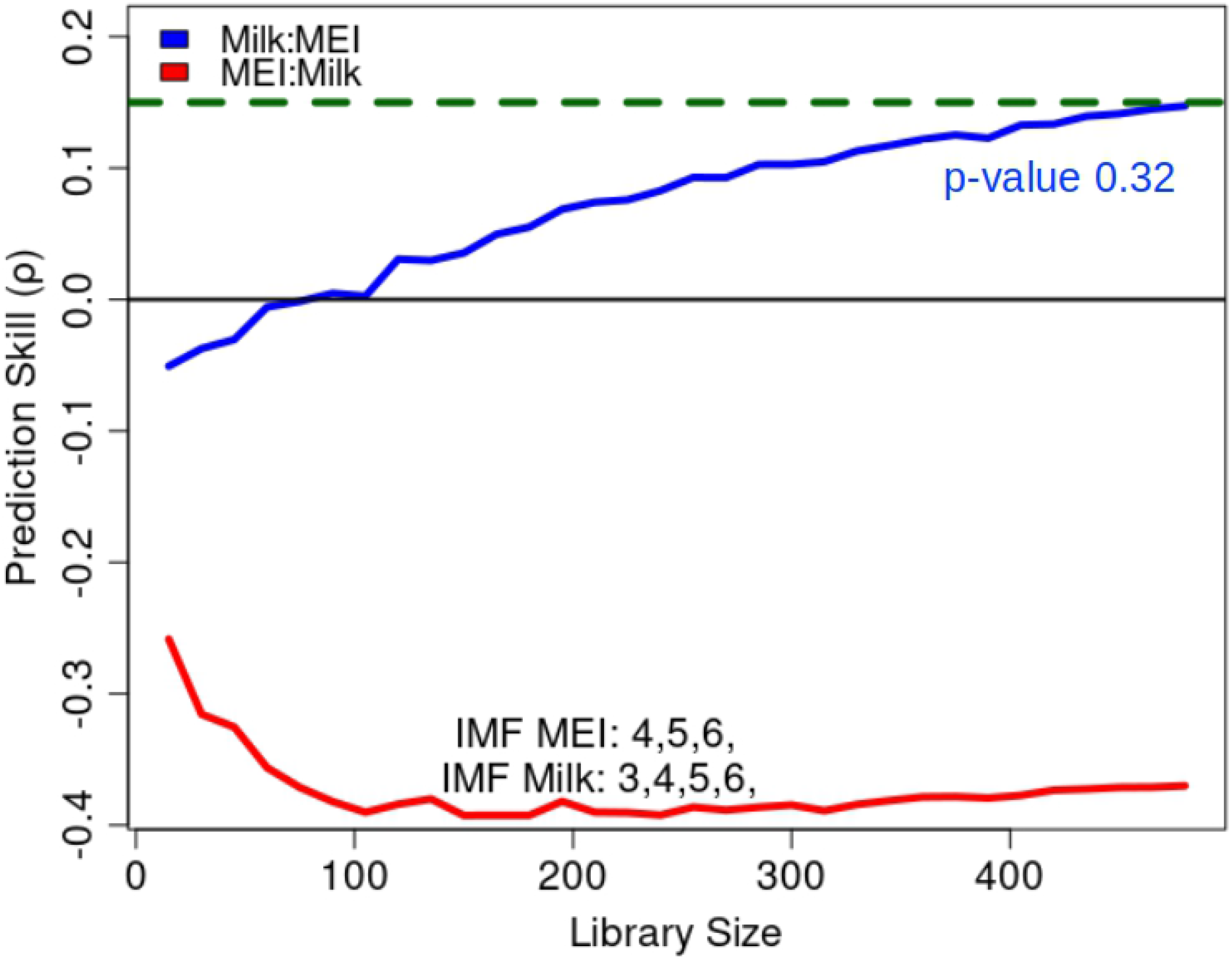
Convergent cross mapping between interannual components of milk production and MEI. Dashed horizontal line is linear cross correlation.

**Fig 8.**
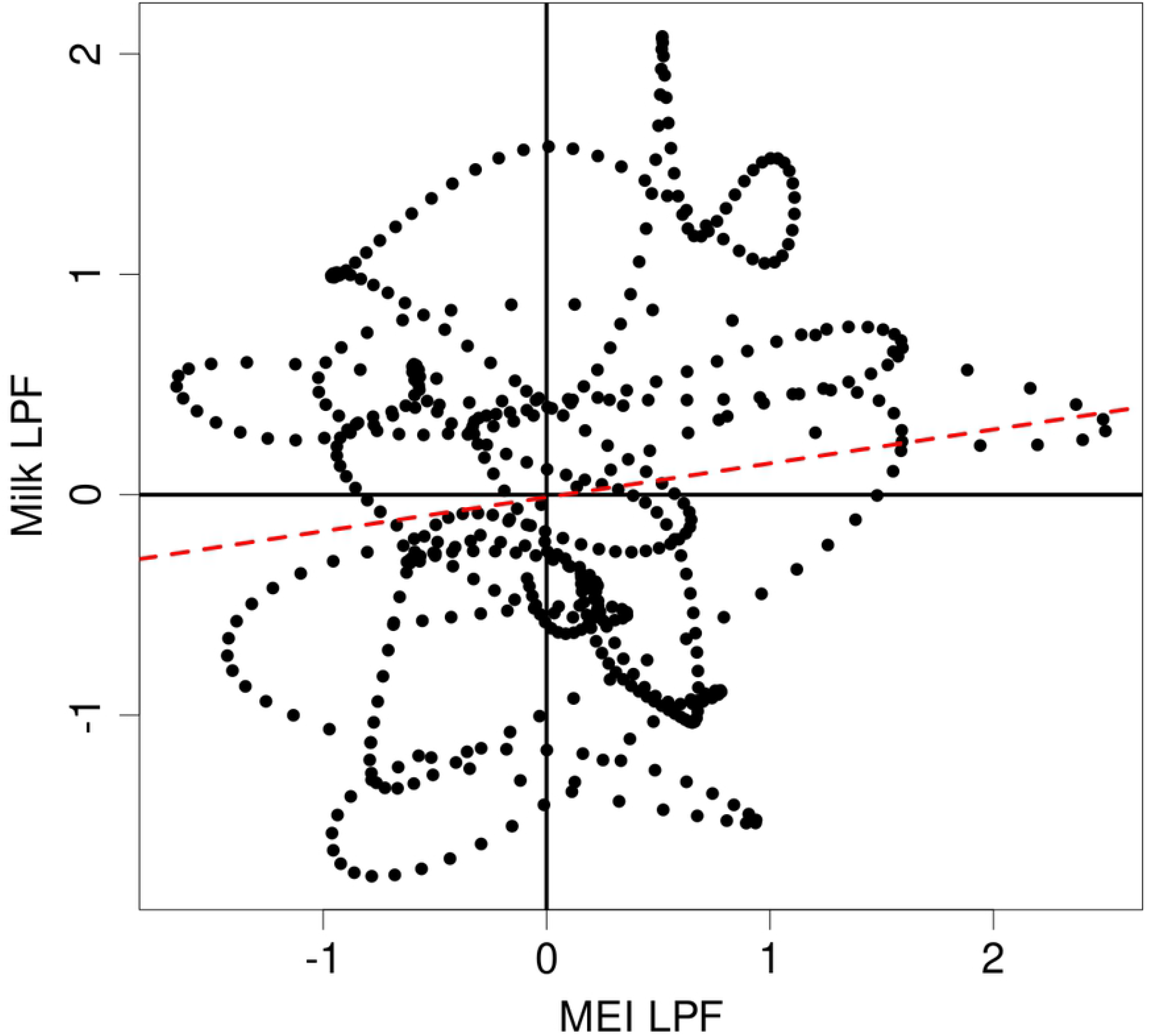
Interannual components of milk production vs. MEI. Dashed red line is the linear regression.

## Acknowledgments

This work was funded in collaboration of the U.S. Department of the Interior, National Park Service, Everglades National Park, and University of California San Diego through the Cooperative Ecosystem Studies Units (CESU) Network http://www.cesu.psu.edu/. This work was supported by DoD-Strategic Environmental Research and Development Program 15 RC-2509, NSF DEB-1655203, NSF ABI-1667584, DOI USDI-NPS P20AC00527, NSF-IOS 1936674, the Scripps Institution of Oceanography Postdoctoral Fellowship, the McQuown Fund and the McQuown Chair in Natural Sciences, University of California, San Diego.

